# Comprehensive Profiling of Monkeypox Virus Antigens Identifies Potent Targets for Next-Generation mRNA Vaccine Development

**DOI:** 10.64898/2026.06.23.733206

**Authors:** Alexandra C. Walls, Harman Malhi, Gavin M. Palowitch, Charles L. Dulberger, Prutha Tarte, Meghan Marquette, Sam Hurbines, Alibek Galeev, Hannah A. Miller, Ehsan Mehravar, Hossam Hefesha, Richard B. Gaynor, Asaf Poran, Adam Zuiani

**Affiliations:** BioNTech US Inc., Cambridge, MA, 02139, USA; Beacon Hill Life Sciences, Boston, MA, 02108, USA; BioNTech SE, 55131 Mainz, Germany

**Author notes:** corresponding author and lead contact: Adam Zuiani, 75 Sidney St, Cambridge, MA 02139 United States. corresponding author: Asaf Poran, 75 Sidney St, Cambridge, MA 02139 United States. equal contribution (Alexandra C. Walls, Harman Malhi, Gavin M. Palowitch, Charles L. Dulberger).

**Keywords:** mRNA vaccine, monkeypox virus, vaccinology, immunology, O*rthopoxvirus*, pandemic preparedness

## Abstract

The 2022 Monkeypox virus (MPXV) outbreak renewed interest in vaccines for orthopoxviruses. Initial development efforts focused on well-established antigen targets, especially A35, B6, and M1. However, orthopoxvirus surfaces are complex, displaying many antigens across two infectious forms, mature virions (MV) and extracellular virions (EV) and targets relevant to protection remain to be comprehensively defined. We leveraged advances in orthopoxvirus protein biology and mRNA vaccine technology to compare immunity to all feasible targets. Mice were immunized with mRNAs encoding each antigen, or antigen complex, and neutralizing antibody responses were measured prior to heterologous challenge with vaccinia virus. Among MV antigens, A28 induced potent complement-mediated neutralizing antibodies, and the A17:G10 complex induced neutralizing antibodies and protected from challenge. For EV antigens, A36 induced neutralizing antibodies and protected from challenge. Our results affirm the consensus strategy focusing on key antigens while highlighting additional targets that could enhance updated MPXV mRNA vaccines.

**SIGNIFICANCE:** Monkeypox virus, a member of the *Orthopoxvirus* genus along with variola virus, has been associated with two recent outbreaks of mpox disease leading to a renewed focus on orthopoxvirus vaccine development. We report an agnostic screen of all monkeypox virus surface antigens where we combined recent advances in structural biology and mRNA technology to evaluate these potential new vaccine targets. We confirmed that historically prioritized antigens M1, A35 and B6 were protective but also discovered new antigens of interest including A28, the A17:G10 complex and A36 that can be the targets of protective immune responses. These findings are critical to inform next-generation vaccine designs should novel orthopoxviruses emerge as human pathogens.

## MAIN

Monkeypox virus (MPXV) is an enveloped DNA virus of the *Orthopoxvirus* genus capable of causing mpox disease in humans. Infection can be lethal, with case fatality rates reported of up to 10%.^1,2^ Orthopoxviruses have high conservation in the genes encoding structural proteins, a feature that was successfully exploited to eradicate smallpox through vaccination against variola virus (VARV), the causative agent of smallpox, using vaccinia virus (VACV) as live vaccine.^3,4^ However, decline in population-level orthopoxvirus immunity following cessation of smallpox vaccination has been associated with an increase in mpox outbreaks, most recently in 2022 and 2024.^2,5,6^ These Public Health Emergencies of International Concern (PHEIC) were notable for the emergence of novel strains and first widespread human-to-human transmission.^6,7^ Currently available VACV-based vaccines were insufficient to combat these outbreaks and there is an unmet need for rapidly scalable alternative vaccines against Orthopoxviruses.^8–13^ mRNA vaccine technology is well-suited to meet this need, with potential for rapid scalability demonstrated during the COVID-19 pandemic.^14^ The utility of mRNA vaccines to combat mpox are under evaluation, including several pre-clinical studies^15–30^ and two investigational mRNA MPXV vaccines (BioNTech’s BNT166a and Moderna’s mRNA-1769) have entered clinical trials.^31–34^ The relatively high number of surface antigens displayed on orthopoxvirus virions makes antigen selection for mRNA vaccines targeting *Orthopoxvirus* species more complex than for viral genera such as *Coronaviridae*. While in theory mRNA vaccines could include many or all of these proteins, in practice a few key antigens must be selected to ensure feasibility. This constraint on mRNA vaccines is an important distinction from VACV-based vaccines that deliver the entire viral proteome simultaneously. Additionally, orthopoxvirus replication produces two unique infectious forms, mature virions (MV) and extracellular virions (EV), with distinct antigens on their respective surfaces.^35^ MV are assembled in the cytosol and are not actively released from living host cells. EV form as MV particles, transit through the host secretory pathway and acquire an additional membrane layer; the final EV particle has a double membrane layer, displaying a different set of membrane proteins than the surface of MV particles. Optimal immunity requires responses to both MV and EV antigens and orthopoxvirus mRNA vaccines must therefore encode representatives of both forms.^36–38^ While both the EV and MV membranes have many proteins displayed on their surfaces, current investigational mpox mRNA vaccines have focused on a relatively small subset of these targets. Specifically, MPXV EV antigens A35 and B6 and MV antigens M1, A29, H3 and E8, are frequently selected for inclusion in mRNA vaccine candidates. The orthologs of these MPXV proteins have historically been the focus of subunit, nucleic acid or viral vectored vaccine development against orthopoxviruses species and, especially for A35, B6 and M1, had strong pre-clinical support for their protective capacity at the time of the 2022 MPXV outbreak.^36,37,39–58^ However, it has yet to be determined if the inclusion of additional antigens could augment the response elicited by vaccines.

We previously reported our pre-clinical evaluation of BNT166a, an investigational quadrivalent mRNA vaccine against MPXV which includes the well-established targets A35, B6, M1 and H3.^31^ Recent advances in the study of orthopoxvirus antigens, protein structural modeling, and the flexibility and potency of mRNA vaccines have enabled the rapid study of many antigens simultaneously. Here, we leverage these advances to design mRNA vaccines encoding each feasible EV and MV antigen and compare the ability of each antigen or antigen complex to induce neutralizing antibodies and protect mice from orthopoxvirus disease. Our results affirm the consensus strategy of focusing on the key antigens A35, B6, and M1 to generate immunity to orthopoxviruses but also identify the A17 and G10 complex (A17:G10), A28 and A36 as potential targets in updated mpox mRNA vaccines or in vaccines against future emerging orthopoxviruses.

## RESULTS

### Minimal modifications to MV antigens facilitate localization to the cell surface

Prior to designing mRNA vaccine constructs, we first compiled a list of antigens of interest on the MV and EV surfaces (Supplementary Table 1) and developed protein design strategies for each target. We included all antigens in the MPXV surface proteome with a sufficiently large ectodomain (ECD) to be a plausible target of antibody recognition. No criterion other than ECD size was used for inclusion to achieve an unbiased screen of vaccine targets regardless of proposed antigen function. Our design objective was to generate cell-surface displayed versions of each target in relevant conformations via only minimal modification. Clade Ia MPXV sequences were used as the base for all variant forms. Fully soluble, secreted designs (indicated by “SOL”) were used only when pairing with an integral membrane protein binding partner was expected. Localization of antigen to the plasma membrane was prioritized as it keeps constant the means of B-cell recognition between antigens, consistently allows for the induction of high titer antibody responses in mice, and is commonly used for mRNA vaccines, including for the SARS-CoV-2 spike protein encoded in COVID-19 vaccines including BNT162b2 and mRNA-1273.^59,60^ A summary of constructs evaluated for this work is given in Supplementary Table 2. We evaluated MV and EV antigens separately, beginning with MV proteins (Fig. 1A). We evaluated surface localization of each design through a plasmid-based expression screen in 293T cells. Screening plasmids encoded Flag-tagged or cMyc-tagged versions of each design to allow for detection in the absence of antigen-specific antibody. The Flag and cMyc tags were rationally placed to minimize impact on the overall conformation of the proteins. This plasmid expression strategy (Fig. 1B-C) allowed for rapid down-selection, selecting constructs based on cell surface expression prior to generating lipid nanoparticle (LNP) mRNA constructs.

**Fig. 1:**
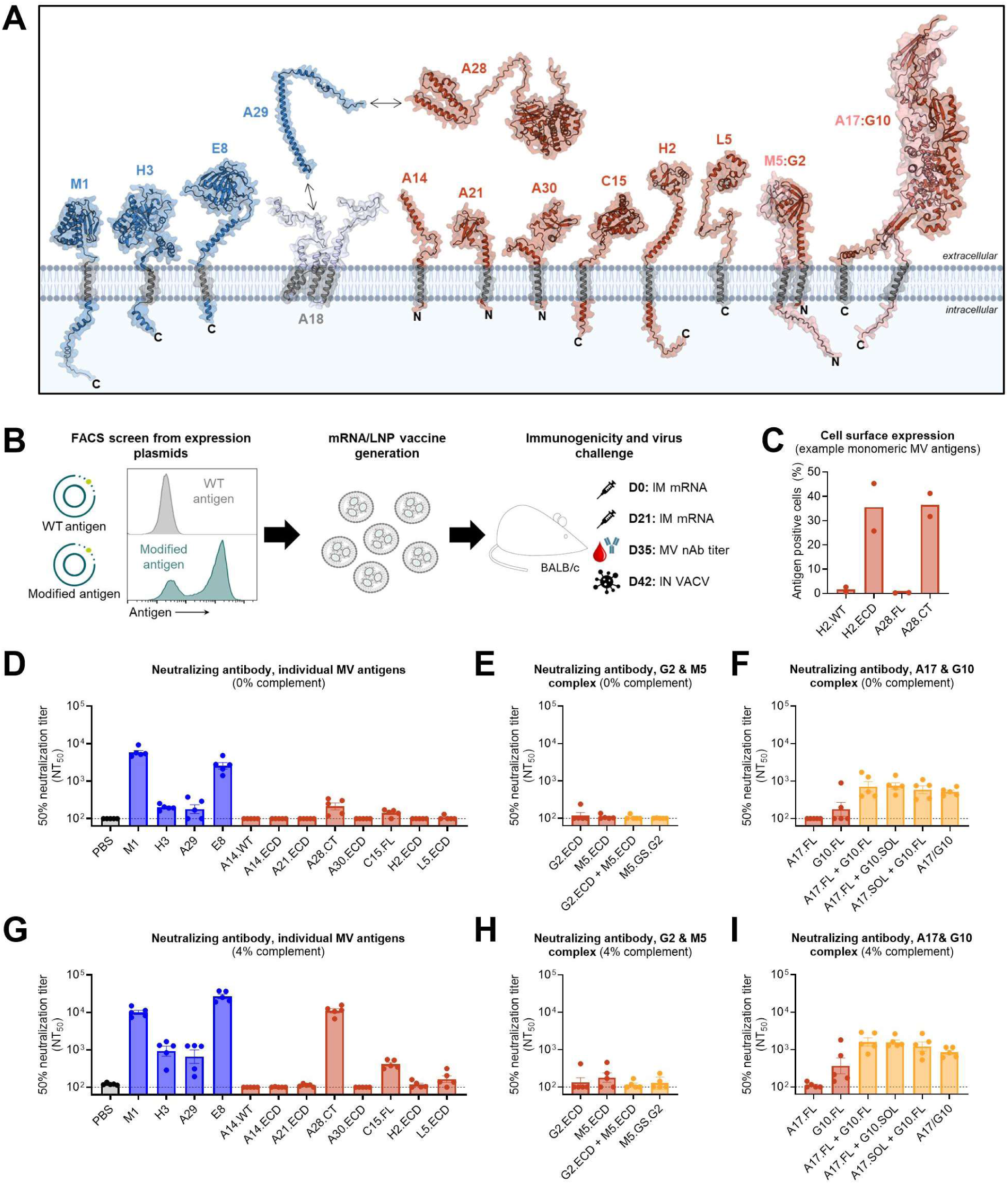
MV antigen evaluation strategy and VACV-neutralizing antibody titers following immunization with optimized vaccine candidates. (A) Schematic displaying all evaluated MV surface antigens. The structures were generated with AlphaFold2 and their accuracy confirmed against X-ray crystallography-derived structures where available (PDB IDs: 1YPY, 6A9S, 4E9O. 5EJ0, 3VOP, 6A9S, 8U0R, 6CJ6, 8INI, 7YTT, 8GP6).^64,65,88–95^ In blue are well-characterized benchmark antigens M1, H3, E8 and A29. In red are novel targets A14, A28, A21, A30, C15, H2, L5, the M5:G2 complex, and the A17:G10 complex. These colors are maintained across the panels, with combination conditions shown in yellow. (B) Visual summary of the antigen design and evaluation process. Different versions of each antigen were encoded on expression plasmids. The plasmids were then used to transfect 293T cells and surface expression was measured by flow cytometry. Designs that facilitated robust expression on the cell surface were prioritized for mRNA vaccine production. Mice were prime-and-boost immunized with these mRNA vaccines, blood collected for measurement of neutralizing antibody responses, and three weeks following the second vaccine dose challenged with VACV-WR IN. (C) Representative flow cytometric surface staining comparing different antigen designs ectopically expressed from plasmids. (D-I) VACV neutralizing titers determined at Day 35 from mice immunized with individual antigens only (D and G) or combinations of antigens known to form heterooligomeric complexes (E, F, H and I). MV-neutralizing antibody was measured in the absence (D-F) or the presence (G-I) of 4% baby rabbit complement. The lower limit of detection is shown as a dotted line, bars represent the geometric mean titer and error bars standard error of the mean (SEM).

A key challenge for ectopic expression of MV proteins relates to the unusual biogenesis of the MV membrane.^61^ MV integral membrane proteins lack signal peptides, are not trafficked to the endoplasmic reticulum and are assembled via mechanisms unique to orthopoxvirus replication.^62,63^ Wildtype (WT) MV protein sequences are therefore ill-suited to ectopic expression in host cells and universally require tailoring to ensure correct display at the cell surface. We attempted different strategies to achieve cell-surface localization depending on the sequence and structure of each MV target. For most targets, multiple designs were evaluated; basic protein schematics and construct names are given in Figure S1A, C & E.

In reviewing MV integral membrane protein sequences, two broad patterns emerged demanding different strategies necessary to ensure display of the ectodomain at the host cell surface. First, many MV targets are arranged as type I integral membrane proteins, with an N-terminal ectodomain and a C-terminal transmembrane (TM) domain but without a signal peptide (SP). These targets (including benchmark antigens M1, H3, E8, and novel targets C15, L5, A17 and G10) could be tailored to cell surface display by simply prepending an SP to the N-terminus. We evaluated whether addition of an HSV-1 gD SP to A17, G10, C15 and L5 alone could result in their trafficking to the cell surface. This signal peptide has previously been used successfully to target MV proteins for translocation to the cell surface.^31^ For C15 and L5, surface expression was evaluated for each construct alone (Figure S1B). For A17 and G10, which form a heterooligomeric complex, various combination transfection conditions were tested (Figure S1F)^64^. Matching expectations, SP-containing full length (indicated by FL) constructs C15.FL and L5.FL were present on the cell surface, however L5.FL expression was low (Figure S1B). We further scrutinized the L5 sequence and observed that L5 has an abnormally short cytoplasmic tail following its TM domain which we hypothesized could compromise ectopic expression. An additional construct, named L5.ECD, in which the endogenous L5 TM was exchanged for an HSV gD TM sequence, was generated and could be readily detected at the cell surface following transfection. Based on these results, C15.FL and L5.ECD were selected as vaccine candidate designs.

We hypothesized that co-expression may be necessary for optimal stability and surface expression of A17 and G10 as these proteins are reported to be paired on the cell surface via an extensive interface. We evaluated surface display of constructs A17.FL and G10.FL, in which an N-terminal SP is prepended to the WT A17 and G10 sequences, respectively, following individual plasmid transfections. While G10.FL could be detected, A17.FL was not detected at the cell surface when expressed alone, but co-transfection of A17.FL with G10.FL rescued A17.FL expression (Figure S1F). We also evaluated an alternative approach to anchoring A17 and G10 to the plasma membrane in which one component was expressed as an integral membrane protein (A17.FL or G10.FL) and the other as a truncated soluble form in which the TM domain was removed (A17.SOL or G10.SOL). A17.SOL and G10.SOL were not detected at the cell surface when expressed alone, an expected outcome given they have no TM domain, but could be detected at the cell surface when co-expressed with G10.FL or A17.FL, respectively. To determine if the complex could be delivered via one mRNA, which would be advantageous in minimizing the number of mRNAs, we also designed a genetic fusion of A17.FL and G10.FL separated by a P2A ribosomal skipping sequence (dubbed A17/G10). A17/G10 expression was measured via a tag on A17, which was readily detected following plasmid transfection. Because of uncertainty about the optimal means of achieving A17 and G10 pairing *in vivo*, several combinations of A17 and G10 constructs were selected for mRNA vaccine production and immunization of mice.

The second pattern we observed among a subset of MV antigens was the presence of a TM domain at the N-terminus with a C-terminal ectodomain. These targets are similar to host type II integral membrane proteins. However, in contrast to host proteins, they had very few hydrophilic residues at their N-termini prior to their TM domains, suggesting they may not integrate efficiently into host membranes in the correct orientation (C-terminus out) with their WT sequence. For these targets (A14, A21, A30, H2, G2 and M5), we designed constructs that display their ectodomains without the endogenous N-terminal TM domain by prepending an artificial SP to the N-terminus of the ectodomain alone and adding a C-terminal artificial TM domain (Figure S1A, dubbed A14.ECD, A21.ECD, A30.ECD, H2.ECD, G2.ECD and M5.ECD). These artificial display approaches were compared to WT versions of each antigen (A14.WT, A21.WT, A30.WT, H2.WT, G2.WT and M5.WT). Expression of G2 and M5 designs were evaluated separately from the other constructs as, similar to A17 and G10, G2 and M5 form a heterodimer and required co-transfection to achieve surface expression.^65^ A14.WT and A30.WT were present on the cell surface at low levels, however A14.ECD, A21.ECD, A30.ECD, and H2.ECD all exhibited enhanced surface expression compared to WT constructs and were selected for mRNA vaccine production (Figure S1B). A14.WT was also advanced to mRNA vaccine production as its expression level was more similar to A14.ECD than that observed for other antigens.

For the G2:M5 complex, in addition to WT and ectodomain-scaffolded variants, we tested soluble ectodomain forms (G2.SOL and M5.SOL) and a fusion protein linking G2 to M5 via a glycine/serine linker (G2.GS.M5) outlined in Figure S1E. G2.WT and M5.WT were both poorly expressed. G2.ECD and M5.ECD could be detected at the cell surface, with G2 expression being relatively weak. No combination of co-expression or fusion of the G2 and M5 complex resulted in increased expression on the cell surface, suggesting that they are not dependent on one another to express, possibly due to their small size and interaction surface. G2.ECD, M5.ECD, and G2.GS.M5 were selected for mRNA vaccine production.

Our final target, A28, was a unique case. A28 has no native TM domain and is thought to anchor to the membrane via a disulfide link to A29, which itself is anchored via an interaction with the integral membrane protein A18.^66,67^ A28 also contains a long aspartate-rich stretch at its C-terminus that we hypothesized may be dispensable for generating immunity and may compromise ectopic expression. We tested two constructs displaying A28 at the plasma cell membrane; one containing the full sequence with cysteine to serine mutations (to prevent aggregation in the absence of binding partner A29) and an artificial SP and C-terminal TM domain (A28.FL), and a second design with only the globular N-terminal component of A28 (A28.CT), eliminating the poly-aspartate sequence (Figure S1A). Only A28.CT successfully expressed on the cell surface (Fig. 1C), suggesting that removal of the aspartate-rich, A29-binding region was required for independent host-cell surface expression and folding. Therefore, A28.CT was selected for mRNA vaccine production.

### A28 and the A17:G10 complex induce MV-neutralizing antibody responses following immunization

To evaluate the mRNA vaccine designs selected from the MV antigen expression screen, we performed combined immunogenicity and protection studies in BALB/c mice. Mice were immunized with individual or combinations of mRNAs, or mock treated with saline on Days 0 and 21 (Fig. 1B). At each immunization day, 1 µg of each mRNA was administered intramuscularly (IM). Serum was collected two weeks following the second vaccination and functional antibody responses evaluated using VACV-neutralization assays. Finally, mice were challenged with a lethal dose of VACV on Day 42.

VACV was selected in lieu of MPXV for neutralization and challenge studies for several reasons. Since our vaccine candidates were designed with MPXV sequences, challenge with VACV arguably represents a more stringent heterologous approach than MPXV challenge. Heterologous protection is of particular importance for orthopoxvirus vaccines which must provide robust cross protection to match historical vaccine performance.

Furthermore, MPXV does not readily infect immunocompetent mouse strains like BALB/c or C57BL/6. MPXV-infection modeling in mice is currently only established in the CAST/Ei strain, which are susceptible to MPXV in part because they are partially immunocompromised.^68^ CAST/Ei mice also demand hybrid EV/MV immunity to achieve protection from disease and therefore are ill-suited to the study of individual antigens, whereas single antigen immunization can protect from VACV.^31^

MV-neutralizing antibody responses induced by the vaccines were measured in the presence and absence of complement. Performance of the MV control antigens M1, A29, H3 and E8, matched expectations from previous vaccine studies (Fig. 1D, G). M1 immunization elicited the highest titer neutralizing antibody response without the addition of complement. E8 immunization also induced antibodies capable of neutralizing MV in the absence of complement, but at lower levels than M1. In the presence of complement, MV-neutralization could also be observed for A29 and H3, and E8 immunized mice had higher titer responses than M1 immunized mice. These results provide benchmarks to evaluate the responses induced by the novel target designs. Most novel antigens tested failed to induce MV-neutralizing antibody responses (Fig. 1D-E, G-H), however, two exceptions were observed. A28 immunization induced no detectable neutralizing antibodies in the absence of complement, but in the presence of complement A28 induced high-titer MV-neutralizing antibody responses, almost matching those observed for M1 and E8. While A17 and G10 individually failed to induce meaningful responses, any combination immunization using the A17:G10 complex induced neutralizing antibody titers both with and without complement (Fig. 1F, I). These data suggest that co-expression may be required for A17 due to the large protein–protein interface demonstrated in the heterodimer complex and may allow these antigens to access more relevant conformations.^64^

### Immunization with the A17:G10 complex matches protection provided by best-in-class MV antigen M1 in orthopoxvirus challenge

Intranasal (IN) VACV-WR challenge in BALB/c is a lethal orthopoxvirus disease model, characterized by rapid weight loss and death in naïve animals. Among benchmark MV antigens, M1 is unique in that immunization against M1 alone can provide complete protection from death and partial protection against weight loss following VACV-WR challenge. In most reports, immunization with other MV antigens fails to provide meaningful survival benefit. In screening novel MV targets we sought to find antigens that could provide survival benefits in the VACV-WR challenge model, ideally matching the performance of M1.

Three weeks following immunization, cohorts of immunized BALB/c mice were challenged with 5 x 10^4^ PFU of VACV-WR IN. Among the benchmark antigens, only M1 provided robust protection (Fig. 2A). E8 immunization resulted in a partial survival benefit, with 30% of animals surviving to the end of the study, while H3 and A29 provided no benefit compared to saline (Fig. 2B-D). Most novel MV antigens provided no benefit with two exceptions. A28 provided a partial survival benefit, with 30% of mice recovering from VACV-WR challenge. While this is modest compared to M1 immunization, it exceeds the protective capacity reported for most other MV antigens and is notable (Fig. 2E-L, Fig. 3A-C). Strikingly, immunization with the A17:G10 complex provided nearly complete protection, mirroring the protection elicited by M1 immunization. Delivery of both components of the complex provided optimal protection, however immunization with G10 alone also provided a degree of protection, with 50% of animals surviving VACV-WR challenge (Fig. 3D-F). Taken together, the neutralization titer and challenge data suggest A28 and A17:G10 may represent previously unappreciated targets of neutralizing antibody activity and protective immunity to orthopoxviruses.

**Fig. 2:**
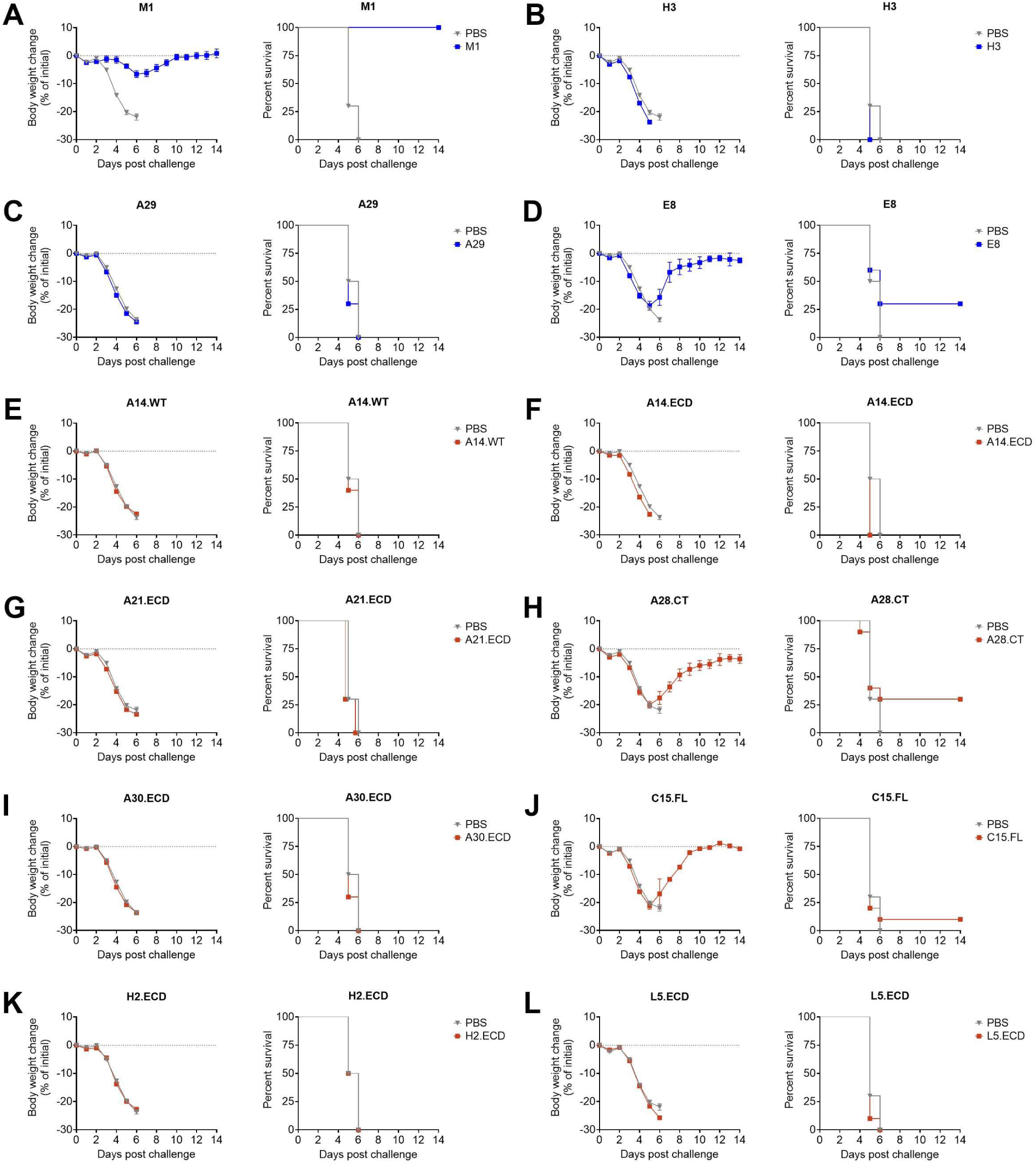
Protection from VACV-WR challenge following immunization with individual MV antigens. **(A-L)** Weight loss (left plots) and survival (right plots) of groups of 10 BALB/c mice immunized with individual MV antigens following IN challenge with VACV-WR. Well-characterized benchmark antigens are shown in blue **(A-D)** and exploratory MV vaccine antigens in red **(E-L)**. Saline control mice are included in each plot in grey. Weight loss is shown as mean percent change in body weight with error bars representing SEM.

**Fig. 3:**
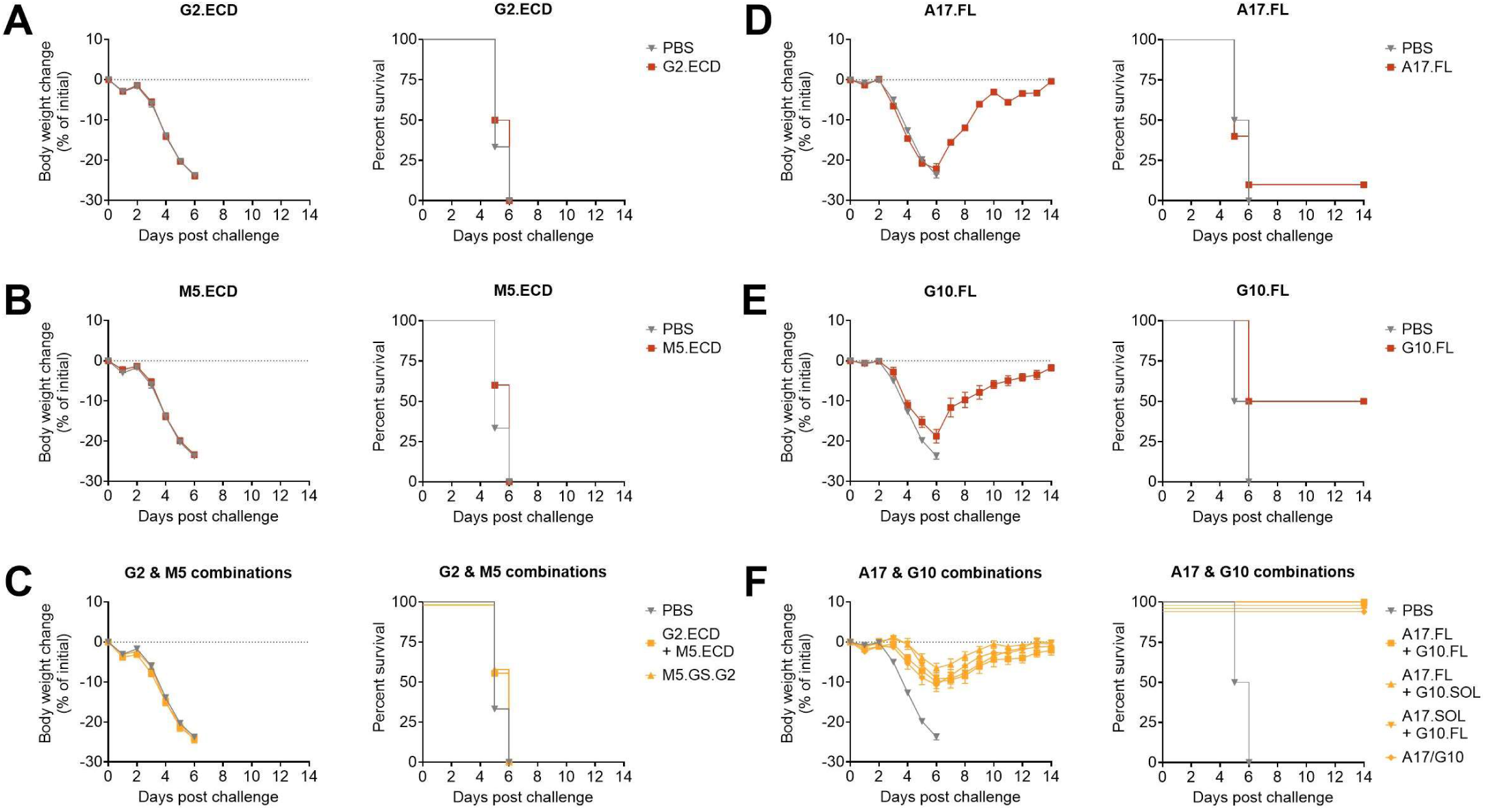
Protection from VACV-WR challenge following immunization with MV antigen complexes. **(A-F)** Weight loss (left plots) and survival (right plots) of groups of 10 mice immunized with MV antigens forming heterooligomeric complexes, G2,M5 and A17,G10, following IN challenge with VACV-WR. Groups immunized with only one component are shown in red **(A, B, D, and E)** and combination immunizations are shown in yellow **(C, F)**. Saline control mice are included in each plot in grey. Weight loss is shown as mean percent change in body weight with error bars representing SEM.

### EV antigens among the B2 complex are detected at the cell surface but A36 must be modified for efficient display

Following our efforts to evaluate MV antigens, we conducted a similar experiment series for EV antigens B2, C2, D14 and A36 (Fig.4A). We evaluated design panels from plasmid-based ectopic expression in 293Ts, generated mRNA vaccines and then measured EV-neutralizing antibody responses and whether immunization with the targets could mitigate orthopoxvirus disease in the VACV-WR model. However, there are relatively few surface proteins on the EV membrane and, in contrast to MV antigens, EV antigens do traffic through the endoplasmic reticulum natively and do not require engineering to their sequences in this regard. We focused on a more limited set of designs using WT sequences for EV proteins (Figure S2A).

**Fig. 4:**
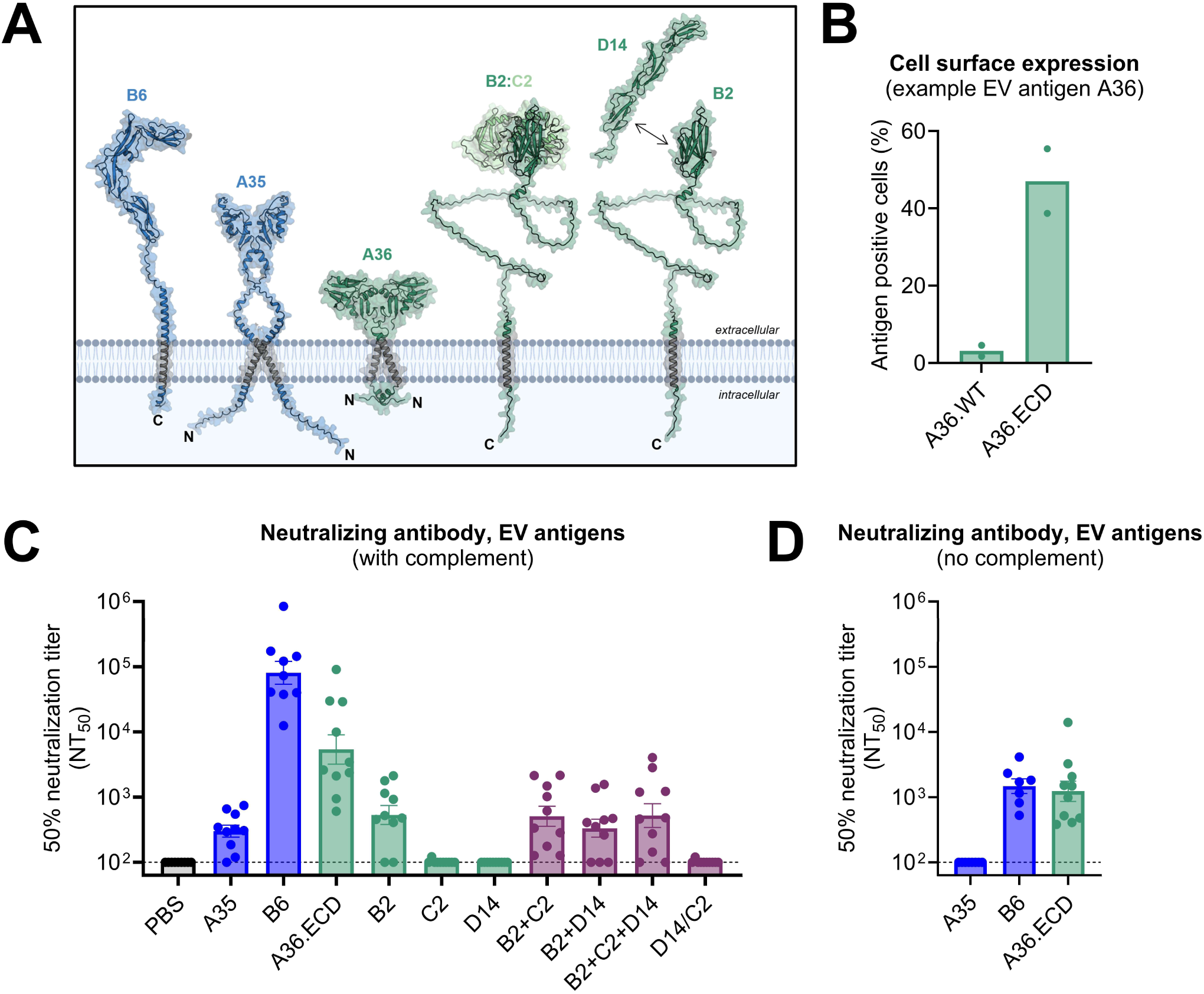
EV antigen evaluation strategy and VACV-neutralizing antibody titers following immunization with EV antigen vaccines. **(A)** Schematic displaying all evaluated EV surface antigens. The structures were generated with AlphaFold2 and their accuracy confirmed against X-ray crystallography-derived structures if available (PDB IDs: 8XS3, 4LQF, 9HL2).^96–98^ In blue are well-characterized benchmark antigens A35 and B6. In green are novel targets A36, B2 in complex with C2, and B2 in complex with D14. These colors are maintained across the panels, with combination conditions shown in purple. **(B)** Representative FACS surface staining comparing different A36 designs ectopically expressed from plasmids. **(C and D)** VACV EV-neutralizing titers determined at Day 35 from mice immunized with individual or combinations of EV antigens EV-neutralizing antibody were measured in the **(C)** presence or **(D)** absence of 4% baby rabbit complement. The lower limit of detection is shown as a dotted line, bars represent the geometric mean titer and error bars are SEM.

We first evaluated expression and assembly of B2 with C2 and D14 at the cell surface. B2 (also referred to as poxvirus hemagglutinin), C2 (also referred to as serine protease inhibitor [SPI]), and D14 (also referred to as virus complement-control protein [VCP]) are EV membrane-associated proteins. B2 is a type I integral membrane protein that forms two heterodimeric complexes with either C2 or D14, which are otherwise soluble proteins.^69–72^ In contrast to MV membrane proteins, each has its own native signal peptide. B2 could be readily detected at the cell surface regardless of whether it was delivered individually or in combination with C2 or D14 (Figure S2B). C2, although it is a secreted, soluble protein, could be detected at the cell surface in all conditions as well, suggesting it can adhere to the surface of 293T cells independent of its interaction with B2. D14 could not be detected at the cell surface when expressed alone, but could be weakly detected if co-expressed with B2, consistent with B2 anchoring D14 to the membrane. We also evaluated a strategy to display C2 and D14 on the plasma membrane without B2 by generating a D14/C2 fusion protein anchored with an artificial TM domain (Figure S2A-B). Consistent with D14 expression patterns, D14/C2 could only weakly be detected at the cell surface following ectopic expression.

A36 is a type II integral membrane protein with a C-terminal ectodomain (Figure S2A) that is proposed to interact with A35 and B6.^73–75^ We initially characterized only the WT sequence (A36.WT) and found very low levels of protein at the cell surface. It is possible that co-expression with A35 and/or B6 could rescue A36, but this would render an evaluation of its individual potential as a target of immune responses impossible as immunization with either A35 or B6 alone induces EV-neutralizing antibodies and fully protects against VACV-WR. We therefore evaluated whether we could display the A36 ECD at the plasma-membrane using an artificial TM domain (A36.ECD). This construct was robustly displayed at the cells surface (Figure S2C) and was selected for use as a vaccine sequence.

### A36 immunization induces potent EV-neutralizing antibody responses

As for MV antigens, once we had selected antigen designs based on *in vitro* expression screening, we proceeded to combined immunogenicity and challenge studies in BALB/c mice following the same experiment design but using EV neutralization assays (Fig. 1B). A35 and B6 are the two cardinal EV antigens found in most orthopoxvirus subunit and mRNA vaccine designs, including BNT166a, and serve as benchmark EV antigens. Both targets are known to induce antibodies capable of neutralizing EV particles in the presence of complement, with antibodies raised to A35 also capable of inhibiting spread of EVs between infected cells.^76,77^ Of note, complement is reported to be a strict requirement in EV neutralization assays, in contrast to MV particles.^78,79^ We first evaluated whether all constructs could generate EV-neutralizing antibody titers in the presence of complement (Fig. 4C) then proceeded to determine if select designs could neutralize EV absent complement (Fig. 4D).

Per expectation, immunization with both A35 and B6 induced antibodies capable of neutralizing EV in the presence of complement, with B6 vaccination generating especially high titer responses. Immunization with A36.ECD also generated EV-neutralizing antibodies, yielding intermediate titers between A35 and B6 in magnitude. Immunization with B2 either alone or in combination with C2 or D14 induced measurable EV-neutralizing antibody titers with complement, but immunization with C2 or D14 individually or as a fusion protein did not produce antibodies capable of neutralizing EV. Interestingly, antibodies raised to B6 and A36, but not A35, could neutralize EV absent complement.

### A36 immunization fully protects from VACV-WR infection and responses raised to D14 partially mitigate disease

Immunization with either of the benchmark EV antigens, A35 or B6, was sufficient to fully mitigate weight loss and death in the VACV-WR challenge model (Fig. 5A & B). We sought to survey the remaining EV antigens to determine if they could provide a survival benefit either alone or in complex with one natural binding partner. Immunization with A36.ECD achieved this goal, providing complete weight loss and survival benefit (Fig. 5C). Immunization with B2 or C2 alone provided no survival benefit (Fig. 5D & E), whereas 30% of mice immunized with D14 alone were protected (Fig. 5F). A modest 20% survival of challenged mice was observed for a combination immunization with B2 and C2 (Fig. 5G), however combination immunization with B2 and D14 resulted in 90% survival, albeit with modest weight loss observed (Fig. 5H). Immunization with B2, C2, and D14 simultaneously resulted in 50% survival, suggesting inferiority to B2 and D14 alone (Fig. 5I). Immunization with a C2/D14 fusion protein yielded 70% survival benefit (Fig. 5J). These results suggest that D14 is likely the best target for survival amongst this complex.

**Fig. 5:**
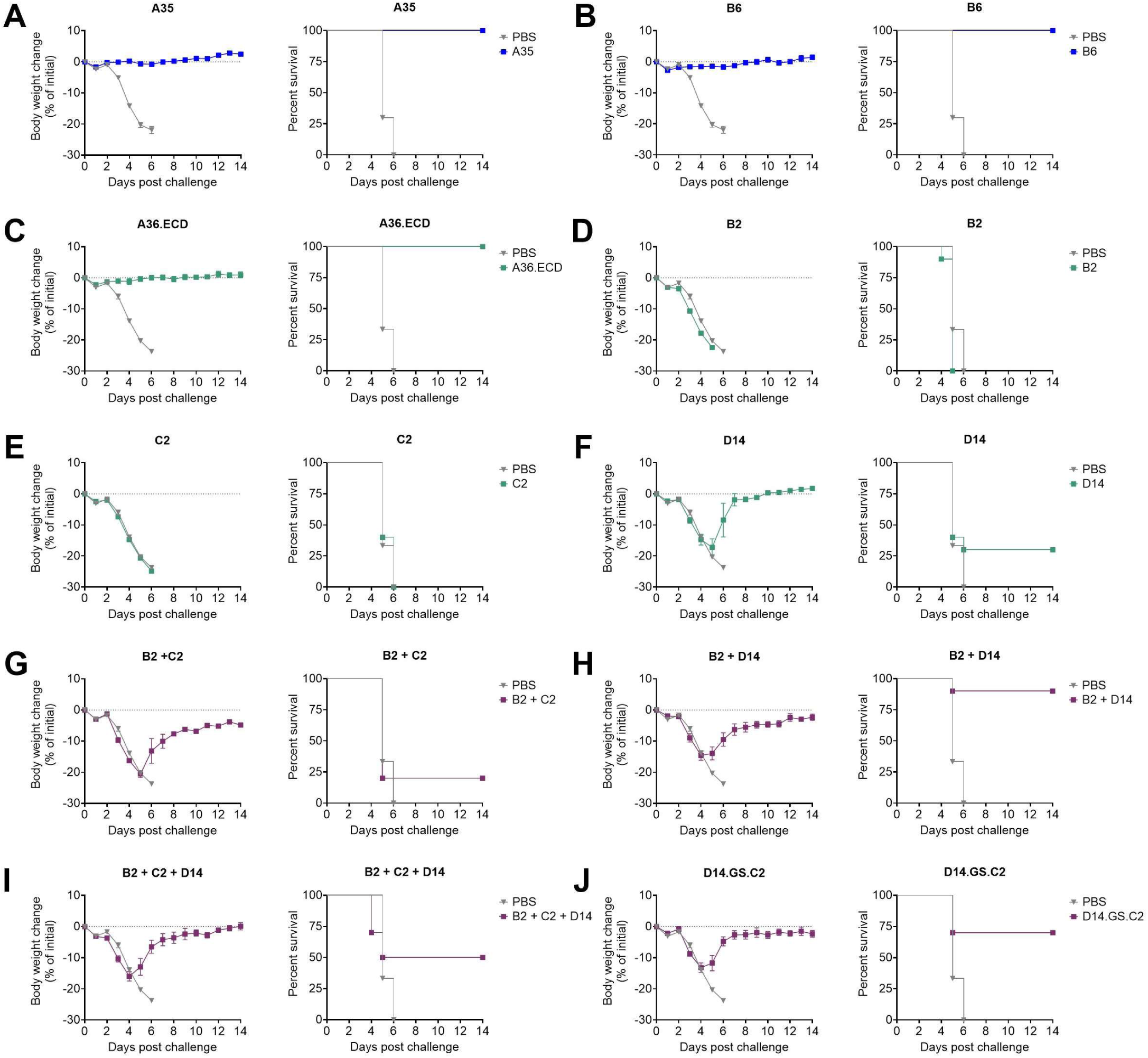
Protection from VACV-WR challenge following EV antigen vaccination. **(A-J)** Weight loss (left plots) and survival (right plots) of groups of 10 mice immunized with EV antigens following IN challenge with VACV-WR. Groups immunized with well-characterized benchmark antigens A35 and B6 are shown in blue **(A and B).** Group immunized with individual exploratory antigens A36, B2, C2 and D14 are given in green **(C-F)** and combination immunizations with B2 complexes members are shown in purple **(G-J)**. Saline control mice are included in each plot in grey. Weight loss is shown as mean percent change in body weight with error bars representing SEM.

### Evaluating antibody responses to surface targets following MPXV infection

Following identification of A36 and A17:G10 as drivers of heterologous protection from VACV-WR challenge, we set out to identify whether antibody responses following MPXV infection in humans would target these antigens. To explore this, we transfected 293T cells with mRNA constructs for each antigen and incubated the transfected cells with serum from mpox convalescent donors prior to running flow cytometric analysis to identify the binding antibody responses. The human sera generally displayed weak or undetectable antibody responses to most surface antigens, including historically explored and protective antigens including M1 or B6 (Fig. 6). Serum antibody responses were detected for some historical benchmarks including A35 or E8. High titers were detected against the EV target B2 across multiple donors (Fig. 6B). These naturally occurring human antibody responses do not correlate with the antigens generating neutralizing or protective antibody responses in the context of VACV-WR challenge. The weak antibody response to key mediators of protection such as B6, M1, A28, A36 or A17:G10 following infection suggests that subunit or mRNA vaccines could exceed the immune responses generated by natural infection by focusing immunity on more effective targets.

**Fig. 6:**
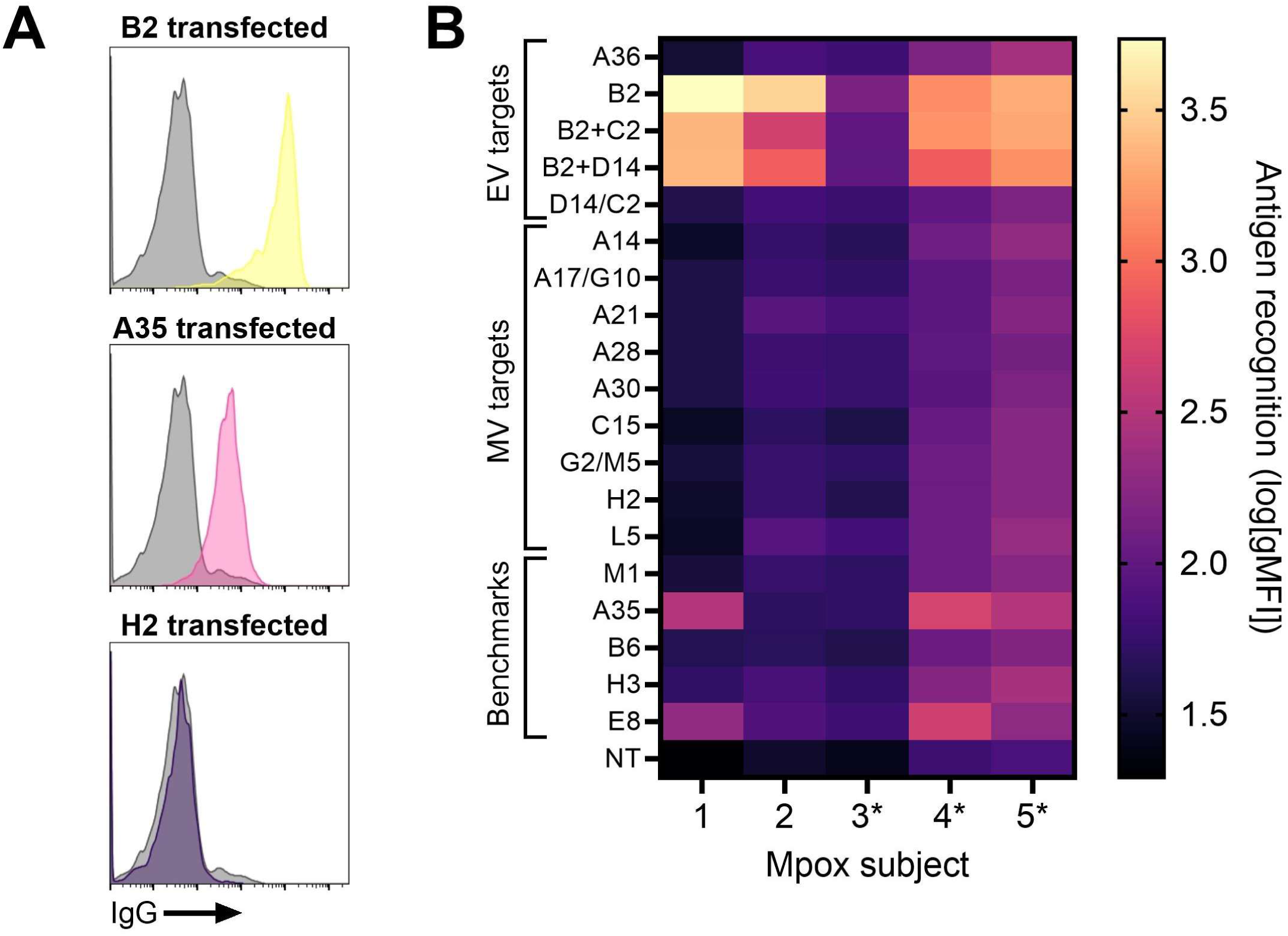
Recognition of mRNA vaccine antigens by mpox convalescent immune sera. **(A)** Representative FACS histograms showing recognition of B2 (yellow, top), A35 (pink, middle) and H2 (purple, bottom) on mRNA vaccine-transfected 293T cells by serum IgG from an mpox convalescent human sera. Staining of non-transfected (NT) control cells is shown in grey in each plot. **(B)** Heat map illustrating the degree of recognition of MPXV antigens by five mpox convalescent individuals. An asterisk denotes individuals with both mpox disease and MVA-BN vaccination.

## DISCUSSION

Successful immunization against smallpox and mpox has employed VACV variants as whole virion vaccines. These vaccines present the natural form and assembly of all viral proteins simultaneously, without modification for ectopic expression in host cells. However, it is unclear whether delivering a high number of targets will necessarily generate an optimal immune response. A focused response to key targets could yield more potent protection against disease. Our comprehensive functional screen of MPXV surface proteins reaffirmed the importance of A35, B6, and M1 for orthopoxvirus immunity and identified additional targets – A36, A28, and the A17:G10 complex – that hold promise for inclusion in future mRNA vaccines. Notably, immune responses raised to most other targets were not individually protective or beneficial, contributing little to neutralizing antibody responses and failing to protect mice from viral challenge. Together, these results show that broad immunity is not necessarily advantageous for protection but expanding beyond A35, B6 and M1 to include additional targets could enhance protection against MPXV and other orthopoxvirus species. Future non-human primate and human vaccine studies to explore the potential benefits of higher valency mpox mRNA vaccines are warranted. Furthermore, our investigation of immune responses in human mpox convalescent sera highlight a potential limitation to relying on natural immunity elicited by orthopoxvirus infection. Antibodies induced by MPXV infection, in sera from individuals with or without prior MVA-BN immunization, were biased towards targets not predicted to provide optimal protection, for instance the B2 complex or E8, while no or weak responses were detected to protective targets like M1, A36, or A17:G10. Our findings suggest that we can leverage the logistical advantages of mRNA vaccines to enable rapid responses to orthopoxvirus outbreaks without compromising protective immunity by focusing on select antigens.

In conducting our antigen design screen, the two orthopoxvirus virion forms presented distinct design challenges. Antigens on the MV surface assemble in atypical ways and their WT sequences are ill-suited for use in mRNA vaccines.^62,63^ At minimum, MV antigen sequences required the addition of an artificial SP for efficient translocation to the cell surface.^80,81^ For many constructs a full artificial design strategy was needed to ensure the display of the ectodomain on host cell surfaces. Additionally, in two target-specific cases, minor sequence modifications rescued antigen expression and display. Specifically, the native A14 TM needed to be exchanged for an alternative sequence, and the C-terminal, aspartate-rich region of A28 required truncation, narrowing the A28 sequence to the predicted globular region of the protein. Finally, for the A17:G10 complex, co-expression of both targets was crucial for display of A17 at the cell surface. Among EV antigens, relatively few modifications were necessary for B2 and its binding partners C2 and D14, although D14 expression was low regardless of modification. One exception to the use of WT sequences for EV targets was A36; the WT A36 sequence was not efficiently displayed at the cell surface. This result could be explained by a dependence on co-expression of other viral factors for A36 trafficking or stability, possibly A35 and B6. We therefore engineered a version of the protein (A36.ECD) that artificially and efficiently displayed the ectodomain on the cell surface. In summary, judicious modifications to antigens allowed us to fully evaluate the potential of each target, including targets that may have remained unappreciated if WT sequences were used.

M1 is consistently reported as the most effective target in the MV membrane.^31,32^ Immunization with M1 induces high titer neutralizing antibody responses that can be enhanced by, but do not depend on, complement.^31^ Furthermore, immunization with M1 is often reported to be unique in its ability to provide full protection from orthopoxvirus disease when compared to other MV antigens. Here, we compare M1 to a broad array of alternative MV targets and confirm its unique potency as a vaccine target, both in terms of induction of MV-neutralizing antibody and survival benefit following VACV-WR challenge. However, we identify the A17:G10 complex as a similarly valuable target for mRNA vaccines, consistent with a recent report using protein-based vaccination.^82^ Co-immunization of A17 and G10 elicited moderate neutralizing responses stronger than either component alone and the complex fully protected mice from VACV-WR challenge. We also identified A28 as a novel target of interest. Immunization with our tailored A28 design induced neutralization in the presence of complement and provided partial protection from VACV-WR, mirroring the benchmark MV antigen E8.

Interestingly, while there appears to be a correlation between the induction of MV-neutralizing antibody and protection in the VACV-WR model, the relationship is imperfect. M1 elicits the strongest neutralizing antibody response and is highly protective, however E8 induces similar or greater neutralizing antibody titers as A17:G10 with markedly reduced ability to protect mice against viral challenge. Future studies to elucidate the mechanisms of protection for MV-based vaccines should aim to disentangle neutralizing antibody responses, T-cell responses, and other mechanisms of antibody-mediated protection.

Immunization with either benchmark EV antigen, A35 or B6, is consistently reported to provide protection from VACV-WR challenge and can induce EV-neutralizing antibody responses.^31,32,36,37,58^ Additionally, antibodies raised to A35 are reported to inhibit viral spread between infected and non-infected cells.^76,77^ In this study we demonstrate that A36 is a similarly potent target for protection from disease and induction of EV-neutralizing antibody responses. The *Orthopoxvirus* genus is divided into two clades, the North American (new world) and African/Eurasian (old world) species.^83^ A35 and B6 are well conserved across Eurasian/African *Orthopoxvirus* species, however their orthologs in North American species like raccoonpox virus and volepox virus exhibit considerable divergence (Supplementary Table 1). A36 is distinguished by its relatively high conservation in the North American species, making it a potentially valuable addition to novel pan-orthopoxvirus vaccines supporting preparedness to emerging orthopoxvirus outbreaks. Our data suggest immunity raised to B2 and its partners C2 and D14 may also be beneficial. For B2, C2, and D14, each antigen individually provided no or little survival benefit. To observe their full potential benefits, they must be co-delivered, presumably to ensure stable expression and correct folding. Among B2, C2 and D14, D14 appears to be the target of greatest potential impact for immune responses. B2 immunization alone induced EV-neutralizing antibodies but was not sufficient for protection, and D14 alone did not induce neutralizing antibodies but could protect animals from severe disease. These findings suggest D14 provides protection through means other than induction of EV neutralizing antibodies which are yet to be elucidated. D14 antagonizes host complement responses and antibodies raised to D14 may mitigate VACV pathogenesis by disrupting this role in complement inhibition.^84–86^ Alternatively, anti-D14 responses could mediate other antibody effector functions including antibody-dependent cell mediated cytotoxicity, a likely mechanism of protection for anti-EV antibodies, as EV proteins and EV themselves decorate infected cells. Defining the exact mechanisms mediating D14 vaccine protection could further guide antigen optimization efforts. D14 also presented expression challenges, suggesting that further efforts to improve D14 design for enhanced expression may reveal even greater impact in the context of

### VACV-WR animal challenge

Our primary immunological endpoint for this study was the induction of functional antibody responses as humoral immunity is crucial for protection from orthopoxviruses.^87^ However, induction of neutralizing antibody was not requisite for inclusion in the VACV challenge phase in our study design. By advancing mice treated with all vaccine designs to VACV challenge irrespective of their neutralizing antibody response, we remained open to the possibility that T cell or antibody effector functions could mediate protection. Ancillary T cell measurements were planned if protection absent neutralizing antibody induction was observed, however no such cases arose. These results support a cardinal role of functional antibody responses in protection from orthopoxvirus disease.

This study has several limitations. First, given the goal of comprehensive profiling, design of any individual antigen could not be fully explored and optimized. There is the potential that our protein and study design, including our preference for membrane display, limited our capacity to identify all antigens cable of eliciting protective responses. Surveying a wider array of combination immunizations, and optimizing expression, e.g. through amino acid variant screening may further identify a full repertoire of potentially valuable MPXV antigens. Additionally, while the VACV-WR model has been foundational to our understanding of immunity to individual orthopoxvirus antigens, other orthopoxviruses may have different pathogenesis requiring different mechanisms of protection from VACV-WR. Future studies exploring the contribution of these targets to protection from other orthopoxviruses may be beneficial.

In summary, initial efforts to design MPXV mRNA vaccines were correctly focused on the key antigens A35, B6 and M1, but some protective targets may have been overlooked. These excluded antigens may have failed to receive full consideration due to the limited structural information available and historical protein design capabilities. Now, with minimal structural tailoring for ectopic expression, we identify A36, A28, and the A17:G10 complex as potential targets for inclusion in future or updated mRNA vaccines. These targets could increase vaccine potency or breadth to optimize protection and the likelihood that a recombinant protein, vectored or nucleic acid orthopoxvirus vaccine can reach pan-genus protection.

## Data availability

All data reported in this paper may be shared by the lead contact upon reasonable request. Any additional information required to reanalyze the data reported in this paper can be made available from the lead contact upon request.

## ACKNOWLEDGEMENTS

This work was supported in part by funding from the Coalition for Epidemic Preparedness Innovations (CEPI). We are grateful for the support of many of our colleagues at BioNTech, including members of the BNT166 team, especially Dawn Myscofski, Frank Bähner, and Leela Davies, and the Publications team, especially Fiona Powell and Amanda Gallagher. We thank Bhavna Chawla at Bioqual, Inc for her assistance in conducting mouse experiments.

## AUTHOR INFORMATION

### These authors contributed equally: Alexandra C. Walls, Harman Malhi, Gavin M. Palowitch, Charles L. Dulberger

A.C.W., H.M., G.M.P., C.L.D., and A.Z. conceived and designed the study. A.C.W, H.M., P.T., M.M., S.H., A.G., H.A.M., E.M., H.H., and A.Z. developed methods, generated reagents, and carried out laboratory experiments. A.C.W., H.M., G.M.P., C.L.D., P.T., M.M., S.H., A.G., H.A.M., R.B.G., A.P., and A.Z. analyzed the data. A.C.W. and A.Z. wrote the original draft. All authors reviewed and agreed to the final version of the paper.

### Corresponding author and Lead contact

Further information and requests for resources and reagents should be directed to by the lead contact, Adam Zuiani

## ETHICS DECLARATIONS

All authors are current or former employees of BioNTech US Inc. or BioNTech SE, own stock or stock options, and may be inventors on patents and patent applications related to RNA technology and related vaccines. RBG has served on the Board of Directors for Alkermes and Board of Directors for Zai Lab.

## METHODS

### Antigen design screen from plasmid DNA

Plasmids were synthesized by Genscript in the pcDNA3.4 backbone. Plasmids encoded either a flag (DYKDDDDK) or cMyc (EQKLISEEDL) tag in the ectodomain of each protein to facilitate surface detection by flow cytometry. Construct ectodomain boundaries are described in Supplementary Table 1 and clade Ia MPXV sequences were used for all designs. Supplemental Table 2 describes the construct nomenclature and design characteristics used throughout the text. When an exogenous signal peptide or transmembrane sequence is used for scaffolding, it is the HSV gD signal peptide (MGGAAARLGAVILFVVIVGLHGVRG) or the HSV gD transmembrane region (GLIAGAVGGSLLAALVICGIVYWMRRHTQKAPKRIRL). FACS analysis was conducted using 293T cells following plasmid synthesis. Cells were seeded in DMEM supplemented with 10% FBS and 1% PenStrep with a cell density of 120,000 cells/well in a 24-well plate. The cells were incubated overnight at 37°C with 5% CO2 and then transfected with 500 ng of total plasmid using Lipofectamine3000 (ThermoFisher Scientific) according to the manufacturer’s instructions. Transfected cells were prepared for flow cytometry approximately 20 hours following transfection. Culture media was removed and the cells were released from the plate via repetitive pipetting in DPBS. Cells were incubated for 30 minutes at 4 °C with 100 µL of FACS stain solution (Biolegend anti-FLAG-PE, Biolegend anti-cMyc-BV421, and viability dye diluted in DPBS +1% BSA). Cells were washed thrice with DPBS + 1% BSA prior to acquisition with a BD FACSymphony A5 SE.

### Preparation of mRNA-Loaded Lipid Nanoparticles (LNPs)

mRNA-loaded lipid nanoparticles (LNPs) were formulated by combining an aqueous phase containing mRNA (0.2 mg/mL) diluted in 0.05 M citrate buffer (pH 4.0) with an organic phase composed of a lipid mixture dissolved in ethanol. The lipid mixture consisted of BHD-C2C2-PipZ, DSPC, cholesterol, and DMG-PEG2000 at molar fractions of 47.5:40.5:10:2, respectively, with an N/P ratio of 6 and a total lipid concentration of 22.73 mM. The two phases were mixed at a 3:1 volume ratio using a T-mixing junction (0.5 mm ID PEEK) at a total flow rate of 40 mL/min. Syringe pumps (Harvard apparatus, MA, USA) were used to control the flow rate of the solutions. To neutralize and reconstruct the LNP structure, the resulting mixture was dialyzed overnight against 10 mM Tris buffer (pH 7.4) using a Slide-A-Lyser dialysis cassette with a 10K MWCO (Thermo Fisher Scientific, Waltham, MA, USA). The LNPs were then concentrated via ultrafiltration using an Amicon Ultra-15 unit (MWCO 100 kDa) (Merck Millipore Ltd., Carrigtwohill, Co. Cork, Ireland, UFC910096) and subsequently diluted with a sucrose solution prepared in 10 mM Tris buffer and stored at -70±10°C. Sucrose was incorporated as both an isotonic agent and cryoprotectant. The final formulation contained mRNA at a concentration of 0.5 mg/mL, with the buffer composition adjusted to 10 mM Tris and 300 mM sucrose at pH 7.4.

### Mouse immunization and challenge

Studies in mice were conducted at BIOQUAL, INC (Rockville, MD). Housing and care of mice conformed to the standards of the Association for Assessment and Accreditation of Laboratory Animal Care (AAALAC). The Institutional Animal Care and Use Committee (IACUC) of BIOQUAL reviewed and the VACV-WR challenge protocol (protocol numbers 23-054P). Groups of 10 male and female, 5 – 8 week old BALB/cAnNHsd mice (Inotiv) were administered either 1 µg mRNA per construct (2 µg or 3 µg total for combination immunizations) or mock treatment with PBS on Study Days 0 and 21. Three weeks following the second vaccine dose, all animals were challenged IN with 5 × 10^4^ PFU VACV-WR in a total volume of 50 μL per animal (25 µL per nare). Animals were monitored daily for weight loss and survival for 14 days following challenge.

### VACV-RFP MV neutralization protocol

Sera collected two weeks following the second vaccine dose (Study Day 35, n = 5 per group) were analyzed for the presence of MV-neutralizing antibodies. Vero cells were added to 96-well flat bottom plates at a density of 10^4^ cells per well in DMEM supplemented with 10% FBS and incubated overnight at 37°C and 5% CO2. The day following cell plating, VACV-RFP (Imanis OV4007) and serum dilutions were prepared using DMEM supplemented with 10% FBS as diluent, combined either in the presence or absence of 4% baby rabbit complement, and incubated for 1 hour at 37°C. Following this incubation, media was removed from Vero cells and the serum and virus mixtures added to the wells. Plates were incubated overnight at 37°C and 5% CO2 and infected cells were visualized via RFP detection and counted using a Sartorius Incucyte. Percent infection was calculated based on normalizing to the mean infection count observed in wells receiving only virus. NT50 was calculated by non-linear regression in Graphpad Prism.

### EV particle generation and neutralization

EV particles were produced by infecting HeLa with VACV-RFP, VAVC-RFP-EV preparations were maintained at 4°C and used for neutralization assays within two weeks of production. To generate the EV preps, 1.5 x 10^7^ HeLa cells were plated into T175 flasks in DMEM supplemented with 10% FBS incubated at 37°C with 5% CO2 overnight. The next day, cells were confirmed to be approximately 90% confluent, media was removed and infected with VACV-RFP diluted in warm DMEM at an MOI of 0.5. The virus-cell mixture was incubated at 37°C with 5% CO2 for 3 hours. Following incubation, the inoculum was removed and the cells were gently rinsed with warm DMEM. 35 mL of warm DMEM supplemented with 10% FBS was then added to the flask and incubated at 37°C with 5% CO2 until clear CPE developed (about 48 hours). Media was collected and clarified at 450 x g for 10 minutes to remove cells. The EV particle titer was determined via the 50% tissue culture infectious dose (TCID50) method in the presence of 20 µg/mL MV-neutralizing antibody 7D11 to specifically quantify EV. EV neutralization assays were conducted as for MV neutralization assays (serum from Study Day 35, n = 10 per group) but in the presence of 20 µg/mL MV-neutralizing antibody 7D11. Percent infection was calculated based on normalizing to the mean infection count observed in wells receiving only virus. NT50 was calculated by non-linear regression in Graphpad Prism.

### Human serum binding by flow cytometry

293T cells were seeded in DMEM supplemented with 10% FBS and 1% penicillin-streptomycin in 24-well plates at a cell density of 400,000 cells/well 6 hours prior to transfection. For transfection, LNP-formulated RNA was diluted to 400 ng total RNA in 50 μL Opti-MEM for individual constructs, and 200 ng per construct for the combination transfections, and added directly to cells. Plates were gently mixed and centrifuged at 500 × g for 5 min at room temperature before incubating at 37°C and 5% CO2 for 18 hours. Cell culture media was then removed and the cells were released from the plate using repetitive pipetting in DPBS. Heat inactivated human sera from MPXV infected individuals were diluted 25x in DPBS supplemented with 1% FBS and then used to stain the harvested cells. Cells were incubated with human sera for 30 minutes on ice, washed with DPBS + 1% FBS, then stained with anti-human IgG-PE (Biolegend) for 30 minutes on ice. Cells were then washed again three times and analyzed using a BD FACSymphony A5 SE.

