## Supplemental Figures and Tables for "Comprehensive Profiling of Monkeypox Virus Antigens Identifies Potent Targets for Next-Generation mRNA Vaccine Development"

### SUPPLEMENTAL INFORMATION

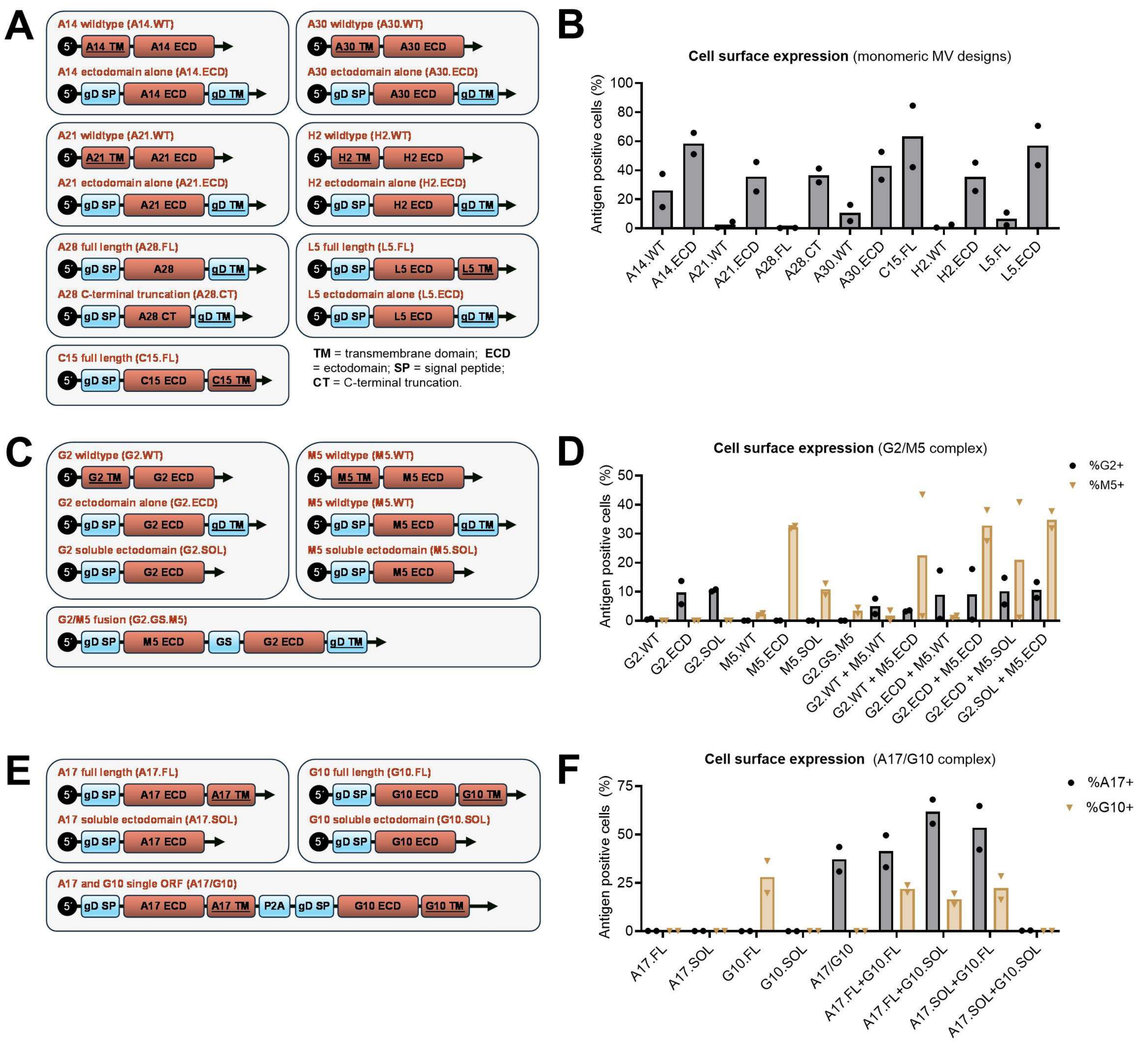


**Figure S1: Surface display of MV antigen designs following ectopic expression form plasmids in 293T cells. (A, C, E)** Schematics representing MV antigen design for **(A)** monomeric or **(C, E)** complex forming proteins. For most targets, several designs were evaluated to determine the minimum modifications necessary to facilitate cell surface expression and complex formation. The sequences of the designs were cloned in pcDNA3.1(+) to allow for ectopic expression in mammalian cells prior to mRNA vaccine production. **(B, D, F)** FACS surface staining comparing WT and engineered antigen designs expressed from plasmids in 293T cells. Tag-based detection was used to eliminate the need for target-specific detection antibodies. For the G2:M5 **(D)** and A17:G10 **(F)** complexes, co-transfections of two plasmids were used to evaluate whether co-expression of both components enhanced detection of each constituent polypeptide. Bars are the mean of two biological replicates.

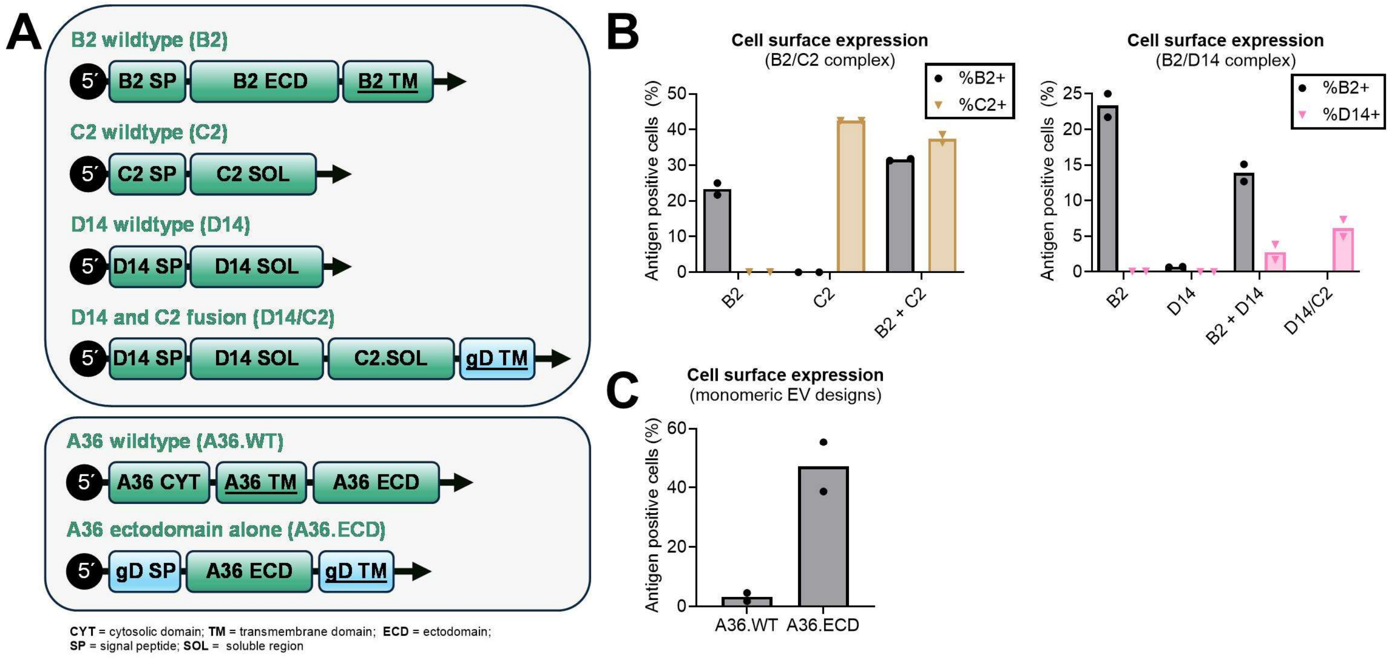


**Figure S2: Surface display of EV antigen designs following ectopic expression form plasmids in 293T cells. (A)** Schematics for WT and engineered EV antigen designs. Two versions of A36 were generated in addition to a fusion of D14 and C2. **(B)** 293T cells were transfected with plasmids encoding WT sequences of B2, C2, and D14, both alone and in combination. Tag-based surface detection was used to eliminate the need for target-specific detection antibodies. Combination transfections of B2 and C2 (left) and B2 and D14 (right) were used to evaluate whether co-expression of both components enhanced detection of each constituent polypeptide. **(C)** 293T cells were transfected with A36 antigen-encoding plasmids and expression at the cells surface evaluated by FACS.

**Supplementary Table 1: MPXV antigen protein characteristics and conservation among orthopoxviruses**

| MPXV name | VACV name | OPG# | Virion form | Length (amino acids) | Size (Kda) | TM domain range | ECD range | VACV Identity | VARV Identity | RCN Identity | VPXV Identity |
|---|---|---|---|---|---|---|---|---|---|---|---|
| A14 | A13 | 139 | MV | 70 | 7.7 | 4 to 23 | 24 to 70 | 93% | 84% | 67% | 60% |
| A17 | A16 | 143 | MV | 377 | 43.7 | 341 to 363 | 1 to 340 | 93% | 96% | 87% | 87% |
| A21 | A21 | 147 | MV | 115 | 13.4 | 4 to 21 | 22 to 115 | 97% | 95% | 88% | 88% |
| A28 | A26 | 153 | MV | 520 | 60.3 | N/A | 1 to 520 | 93% | 90% | 73% | 70% |
| A29 | A27 | 154 | MV | 110 | 12.6 | N/A | 1 to 110 | 94% | 94% | 67% | 66% |
| A30 | A28 | 155 | MV | 146 | 16.4 | 5 to 27 | 28 to 146 | 96% | 97% | 87% | 84% |
| C15 | F9 | 53 | MV | 212 | 23.8 | 177 to 196 | 1 to 176 | 99% | 98% | 92% | 89% |
| E8 | D8 | 120 | MV | 304 | 35.2 | 275 to 294 | 1 to 274 | 95% | 93% | 69% | 68% |
| G2 | G3 | 86 | MV | 111 | 12.8 | 4 to 20 | 20 to 111 | 98% | 95% | 92% | 90% |
| G10 | G9 | 94 | MV | 340 | 38.8 | 322 to 340 | 1 to 322 | 99% | 98% | 88% | 88% |
| H2 | H2 | 107 | MV | 189 | 21.5 | 29 to 48 | 48 to 189 | 99% | 100% | 93% | 92% |
| H3 | H3 | 108 | MV | 324 | 37.5 | 284 to 305 | 1 to 283 | 94% | 94% | 86% | 85% |
| L5 | J5 | 104 | MV | 133 | 15.1 | 110 to 133 | 1 to 109 | 98% | 100% | 89% | 88% |
| M1 | L1 | 95 | MV | 250 | 27.3 | 182 to 204 | 1 to 181 | 99% | 99% | 94% | 95% |
| M5 | L5 | 99 | MV | 128 | 15.0 | 30 to 47 | 48 to 128 | 99% | 99% | 99% | 96% |
| A35 | A33 | 161 | EV | 181 | 20.0 | 34 to 56 | 57 to 181 | 96% | 91% | 73% | 73% |
| A36 | A34 | 162 | EV | 168 | 19.6 | 15 to 38 | 39 to 168 | 95% | 96% | 89% | 92% |
| B2 | A56 | 185 | EV | 313 | 34.4 | 278 to 300 | 19 to 277 | 93% | 80% | 65% | 50% |
| B6 | B5 | 191 | EV | 317 | 35.1 | 280 to 302 | 20 to 279 | 96% | 93% | 66% | 66% |
| C2 | K2 | 40 | EV | 375 | 42.8 | N/A | 16 to 375 | 92% | 92% | 77% | 78% |
| D14 | C3 | 32 | EV | 216 | 23.4 | N/A | 20 to 216 | 96% | 92% | 73% | 69% |

N/A = not applicable; RCN = raccoonpox virus; VPXV = volepox virus

**Supplementary Table 2: Antigen constructs used in expression screen and mRNA vaccines**

| Antigen | Construct name | Abbreviation | Description | mRNA vaccine production |
|---|---|---|---|---|
| A14 | A14 wildtype | A14.WT | Unmodified A14 amino acid sequence | No |
| | A14 ectodomain | A14.ECD | A14 ECD displayed on the cell surface via the addition of an N-terminal HSV gD SP and C-terminal HSV gD TM | Yes |
| A21 | A21 wildtype | A21.WT | Unmodified A21 amino acid sequence | No |
| | A21 ectodomain | A21.ECD | A21 ECD displayed on the cell surface via the addition of an N-terminal HSV gD SP and C-terminal HSV gD TM | Yes |
| A28 | A28 full length | A28.FL | Full A28 sequence displayed on the cell surface via the addition of an N-terminal HSV gD SP and C-terminal HSV gD TM. Two cysteine residues (VACV numbering 441 and 442, MPXV numbering 332 and | No |

| | | | | |
|---|---|---|---|---|
| | | | 333) known to mediate bonding to A29 via disulfides were substituted with serine residues. | |
| | A28 C-terminal truncation | A28.CT | Truncated A28 sequence encoding only the N-terminal globular component displayed on the cell surface via the addition of an N-terminal HSV gD SP and C-terminal HSV gD TM | Yes |
| C15 | C15 full length | C15.FL | Full C15 sequence displayed on the cell surface via the addition of an N-terminal HSV gD SP | Yes |
| A30 | A30 wildtype | A30.WT | Unmodified A30 amino acid sequence | No |
| | A30 ectodomain | A30.ECD | A30 ECD displayed on the cell surface via the addition of an N-terminal HSV gD SP and C-terminal HSV gD TM | Yes |
| H2 | H2 wildtype | H2.WT | Unmodified H2 amino acid sequence | No |
| | H2 ectodomain | H2.ECD | H2 ECD displayed on the cell surface via the addition of an N-terminal HSV gD SP and C-terminal HSV gD TM | Yes |
| L5 | L5 full length | L5.FL | Full L5 sequence displayed on the cell surface via the addition of an N-terminal HSV gD SP | Yes |
| | L5 ectodomain | L5.ECD | L5 ECD displayed on the cell surface via the addition of an N-terminal HSV gD SP and C-terminal HSV gD TM | Yes |
| G2 | G2 wildtype | G2.WT | Unmodified G2 amino acid sequence | No |
| | G2 ectodomain | G2.ECD | G2 ECD displayed on the cell surface via the addition of an N-terminal HSV gD SP and C-terminal HSV gD TM | Yes |
| | G2 soluble ectodomain | G2.SOL | G2 soluble ectodomain secreted from cells via the addition of a N-terminal HSV gD SP | No |
| M5 | M5 wildtype | M5.WT | Unmodified M5 amino acid sequence | No |
| | M5 ectodomain | M5.ECD | M5 ECD displayed on the cell surface via the addition of an N-terminal HSV gD SP and C-terminal HSV gD TM | Yes |
| | M5 soluble ectodomain | M5.SOL | M5 soluble ectodomain secreted from cells via the addition of an N-terminal HSV gD SP | No |
| G2 and M5 | G2/M5 fusion | G2.GS.M5 | A fusion protein encoding an HSV gD SP, M5 ECD, a GS-linker, G2 ECD, and HSV gD TM | Yes |
| A17 | A17 full length | A17.FL | Full A17 sequence displayed on the cell surface via the addition of an N-terminal HSV gD SP | Yes |
| | A17 soluble ectodomain | A17.SOL | A17 soluble ectodomain secreted from cells via the addition of an N-terminal HSV gD SP | Yes |
| G10 | G10 full length | G10.FL | Full G10 sequence displayed on the cell surface via the addition of an N-terminal HSV gD SP | Yes |
| | G10 soluble ectodomain | G10.SOL | G10 soluble ectodomain secreted from cells via the addition of an N-terminal HSV gD SP | Yes |
| A17 and G10 | A17 and G10 single open reading frame | A17/G10 | A single mRNA designed to deliver A17 and G10 simultaneously by encoding an HSV gD SP, A17, a P2A site, a second HSV gD SP, and G10. | Yes |
| B2 | B2 wildtype | B2 | Unmodified B2 amino acid sequence | Yes |
| C2 | C2 wildtype | C2 | Unmodified C2 amino acid sequence | Yes |

| | | | | |
|---|---|---|---|---|
| D14 | D14 wildtype | D14 | Unmodified D14 amino acid sequence | Yes |
| C2 and D14 | D14 and C2 fusion | D14/C2 | A fusion protein encoding D14, a GS-linker, the soluble ectodomain of C2, and an HSV gD TM | Yes |
| A36 | A36 wildtype | A36.WT | Unmodified A36 amino acid sequence | No |
| | A36 ectodomain | A36.ECD | A36 ECD displayed on the cell surface via the addition of an N-terminal HSV gD SP and C-terminal HSV gD TM | Yes |
